# Sulfur amino acid restriction prevents S-adenosylmethionine-driven liver steatosis, hepatocellular carcinoma, and metabolic remodeling in high-fat-fed GNMT-null mice

**DOI:** 10.64898/2026.07.17.738958

**Authors:** Griffin S. Hampton, Andres F. Ortega, Cha Mee Vang, Ferrol I. Rome, Mickael Goelzer, Louise Lantier, Curtis C. Hughey

**Author notes:** Correspondence: Curtis C. Hughey from the Department of Medicine, Division of Molecular Medicine, University of Minnesota, Dwan 316, 425 East River Parkway, Minneapolis, MN, 55455.

## Abstract

The expression of glycine N-methyltransferase (GNMT), a critical regulator of S-adenosylmethionine (SAM) levels, is down-regulated in humans with metabolic dysfunction-associated steatotic liver disease (MASLD) and hepatocellular carcinoma (HCC). In low-fat-fed mice, GNMT knockout (KO) induces liver steatosis that progresses to HCC. This is accompanied by increased SAM and a shunting of tricarboxylic acid (TCA) cycle intermediates away from gluconeogenesis to other biosynthetic pathways that support lipid accretion and tumorigenesis. The objective of this study was to test whether this metabolic remodeling persists in GNMT KO mice with diet-induced obesity and to determine if the liver pathophysiology and metabolic dysregulation are dependent on elevated SAM. To accomplish this, GNMT KO mice and wild-type (WT) littermates were fed a high-fat control or high-fat sulfur amino acid restricted (SAAR) diet to mitigate SAM accumulation. ^2^H/^13^C isotope infusions in mice quantified *in vivo* liver glucose and TCA cycle fluxes. Metabolomics, respirometry, and pyruvate tolerance tests were completed to more fully interpret the ^2^H/^13^C metabolic flux analyses. KO mice had impaired gluconeogenesis sourced from TCA cycle intermediates. A concurrent elevation in metabolites of pathways that use both SAM and TCA cycle intermediates indicated increased liver polyamine turnover, transsulfuration, and *de novo* lipogenesis. Importantly, SAAR prevented the increase in SAM, the associated metabolic dysregulation, and the appearance of liver steatosis and HCC. In conclusion, the results of these experiments suggest that the loss of GNMT in mice with diet-induced obesity rewires metabolism in a SAM-dependent manner that precipitates liver steatosis and the transition to HCC.

**Graphical Abstract:** 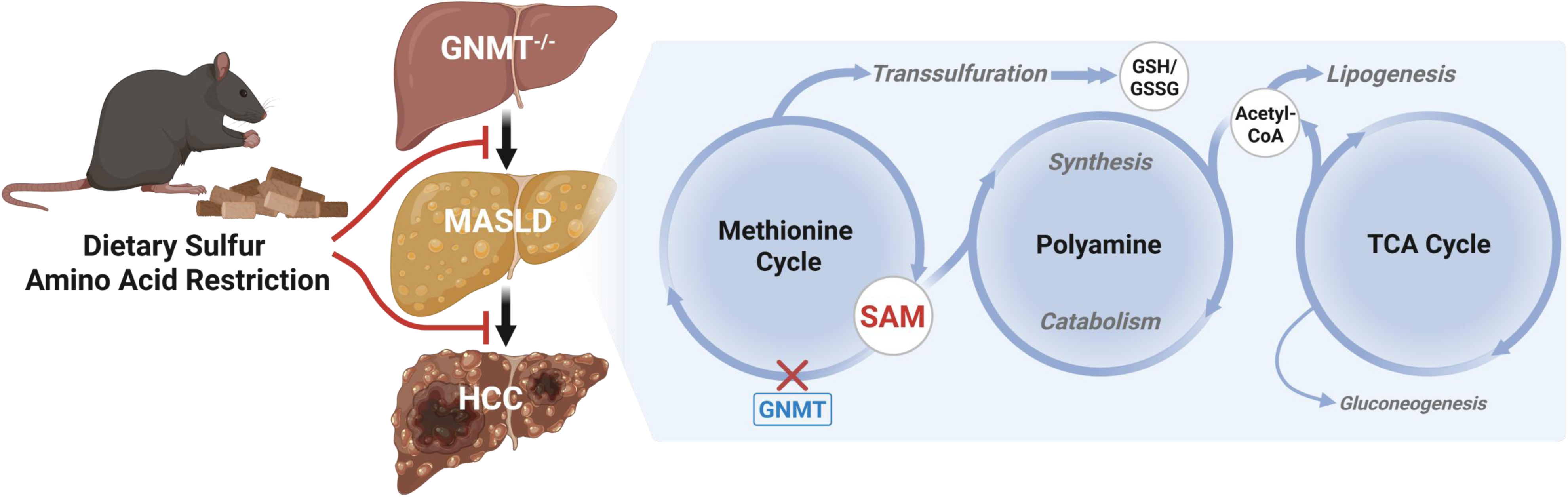

## 1. Introduction

The global prevalence of metabolic dysfunction-associated steatotic liver disease (MASLD) is ∼30% [1]. MASLD is characterized by liver lipid accumulation and is a primary risk factor for advanced liver diseases such as hepatocellular carcinoma (HCC) [2–4]. Although the etiology of MASLD-related HCC is multifactorial, metabolic dysregulation is a key contributor to its pathological progression. In cancer, increased macromolecule synthesis, achieved in part by augmenting intracellular metabolite fate, supports the initiation and maintenance of malignant properties [5, 6]. Dysregulated glucose and mitochondrial metabolism have been implicated as a means by which nutrient fate is rewired across cancer types. In human HCC, glucose production and mitochondrial oxidative metabolism are decreased [7–9]. This decline occurs prior to HCC [10] and is proposed to contribute to an imbalance in anabolic and catabolic pathways, resulting in nutrient flux that supports biomass accumulation [11].

Factors driving metabolic dysregulation in MASLD-related HCC remain unclear. Mouse models of MASLD and HCC are frequently characterized by lower liver glycine N-methyltransferase (GNMT) mRNA and protein levels, which positively correlate with liver injury [12–15]. GNMT is a highly expressed liver methyltransferase that transfers a methyl group from S-adenosylmethionine (SAM) to glycine, forming sarcosine and S-adenosylhomocysteine (SAH) [16]. This enzymatic action is essential for SAM homeostasis, as GNMT knockout (KO) mice exhibit liver SAM concentrations exceeding those of controls by >35-fold [11, 17–20]. A causal role for decreased GNMT in MASLD-related HCC is supported by KO mice, which develop liver steatosis at 12 weeks of age that progresses to HCC by 32 weeks of age [11, 17, 20–23]. The translational significance of this phenotype in mice is underscored by reports of lower or absent GNMT expression in human MASLD [12, 14, 24–26] and HCC [27–29].

Beyond its role as a methyl donor, SAM supports or serves as a substrate for biosynthetic pathways upregulated in MASLD and HCC including lipid, glutathione (GSH), and polyamine synthesis [30–33]. Our prior work in lean mice fed a low-fat diet suggests that GNMT deletion redirects nutrients away from gluconeogenesis and tricarboxylic acid (TCA) cycle oxidation toward these biosynthetic pathways to mitigate elevated SAM [11, 17]. We hypothesize that, due to increased SAM availability, the loss of GNMT promotes a liver metabolic program that supports MASLD-related HCC. Intriguingly, this metabolic remodeling may be diet-dependent. In contrast to the impaired mitochondrial respiratory function observed in low-fat-fed KO mice, isolated liver mitochondria from high-fat-fed GNMT KO mice reportedly display elevated oxidative capacity [34]. Additionally, *in vitro* studies suggest that the tumor suppressor action of GNMT may be independent of its regulation of SAM concentration [35]. It remains unknown whether loss of GNMT *in vivo* promotes HCC independently of SAM.

The objective of this study was to determine if the diminished gluconeogenesis and mitochondrial oxidative metabolism in low-fat-fed GNMT KO mice persist in the presence of diet-induced obesity. We also aimed to test the extent to which metabolic dysregulation, liver steatosis, and HCC in high-fat-fed GNMT KO mice are dependent on increased SAM. GNMT KO mice and wild type (WT) littermates were fed a high-fat control or high-fat sulfur amino acid restricted (SAAR) diet, to prevent SAM accumulation, for 6 or 26 weeks. ^2^H/^13^C isotope infusions in conscious, unrestrained mice were performed to quantify glucose and TCA cycle fluxes *in vivo*. Indirect calorimetry, metabolomics, and molecular analyses were completed to more fully interpret metabolic flux phenotypes. The results show that loss of GNMT impairs gluconeogenesis by blocking TCA cycle cataplerosis and shunting TCA cycle intermediates into *de novo* lipogenesis and polyamine turnover pathways. Moreover, SAAR prevents SAM accumulation, reverses dysregulated nutrient partitioning, and attenuates fatty liver and HCC in KO mice.

## 2. Methods

### 2.1 Mouse model, husbandry, and ethics statement

The University of Minnesota Institutional Animal Care and Use Committee (IACUC) and Vanderbilt University IACUC approved all experimental procedures. Mice with a whole-body KO of GNMT and WT littermates on a C57BL/6J background [11, 17, 18, 20] were used for all studies. Initial experiments testing the interaction between diet-induced obesity and loss of GNMT on liver metabolism and pathophysiology provided WT and KO mice with a high-fat diet (60% kcal from fat; Bio-Serv, F3282, Flemington, NJ) from 6 to 12 weeks of age. Experiments testing the impact of dietary SAAR on liver metabolism and pathophysiology randomly provided WT and KO mice with a high-fat SAA control (0.85% kcal from methionine; 60% kcal from fat; HF-CTRL; A11051306, Research Diets Inc., New Brunswick, NJ) or HF-SAAR (0.125% kcal from methionine; 60% kcal from fat; A11051305, Research Diets Inc., New Brunswick, NJ) diet from 6 to 12 weeks of age when KO mice display fatty liver [17, 20] or from 6 to 32 weeks of age when KO mice develop HCC [11, 21]. A detailed dietary composition for these two diets is provided in Supplementary Table S1. Food and water were provided ad libitum. Mice were housed in temperature (∼22°C)- and humidity (30-70%)-controlled conditions maintained on a 12:12-hour light:dark cycle. Male mice were used for all experiments and were fasted for 8 hours starting within the first hour of the light cycle prior to being euthanized via cervical dislocation. Tissues were then immediately excised, freeze-clamped in liquid nitrogen, and stored at -80°C until analysis.

### 2.2 Body composition and indirect calorimetry

Indirect calorimetry was performed using the Promethion Core System (Sable Systems International, North Las Vegas, NV). Mice were individually housed for six days in temperature (∼22°C)- and humidity (30-70%)-controlled conditions maintained on a 12:12-hour light:dark cycle. The first three days were the acclimation period followed by three days of data collection to assess food intake, locomotor activity, oxygen consumption (VO_2_), CO_2_ production (VCO_2_), respiratory exchange ratio (RER; VCO_2_/VO_2_), and energy expenditure. Energy expenditure was calculated from VO_2_ and VCO_2_ using the Weir equation [36]. Mice used in the indirect calorimetry experiments were 12 weeks of age and had been receiving a high-fat diet (60% kcal from fat; Bio-Serv, F3282, Flemington, NJ) ad libitum for approximately 6 weeks. For mice undergoing indirect calorimetry, body composition was measured using a mq10 nuclear magnetic resonance analyzer (Bruker Corporation, Billerica, MA, USA). In all other cohorts, body composition was determined using an EchoMRI-100™ Body Composition Analyzer (EchoMRI LLC, Houston, TX).

### 2.3 Surgical procedures

At 11 weeks of age, WT and KO mice underwent vascular catheterization surgeries for stable isotope infusion experiments as previously performed [17, 37]. Briefly, carotid artery and jugular vein catheters were implanted for arterial sampling and intravenous infusions, respectively. The free ends of the catheters were exteriorized at the back of the neck, flushed with 5 mg·ml^-1^ ampicillin and 200 U·ml^-1^ heparinized saline, and sealed with stainless-steel plugs. Following surgery, the mice were singly housed and provided approximately seven days of postoperative recovery before stable isotope infusion experiments.

### 2.4 Stable isotope infusions

Access to food and water was withdrawn within an hour of the start of the light cycle. The exteriorized vascular catheters were connected to infusion syringes three hours into the fast. Following an hour acclimation, an 80 μl arterial blood sample was collected to quantify natural isotopic enrichment of plasma glucose. Venous infusions of stable isotopes were performed as previously outlined [11, 17]. In brief, a ^2^H_2_O (99.9%)-saline bolus was administered over a 25-minute period to enrich body water to ∼4.5%. [6,6-^2^H_2_]glucose (99%) was solubilized in the ^2^H_2_O-saline bolus for a prime (440 μmol·kg^-1^). An independent, continuous [6,6-^2^H_2_]glucose (4.4 μmol·kg^-1^·min^-1^) infusion was started after the ^2^H_2_O-saline bolus and [6,6-^2^H_2_]glucose prime. A primed (1.1 mmol·kg^-1^), continuous (0.055 mmol·kg^-1^·min^-1^) intravenous infusion of [^13^C_3_]propionate (99%, sodium salt) was initiated two hours following the ^2^H_2_O bolus and [6,6-^2^H_2_]glucose prime. Four 100 μl arterial samples were collected 90-120 minutes following the [^13^C_3_]propionate bolus (7.5-8 hours of fasting) to determine arterial blood glucose using an Accu-Chek® glucometer (Roche Diagnostics, Indianapolis, IN) and to complete ^2^H/^13^C metabolic flux analysis. Donor red blood cells were resuspended in 4.5% ^2^H_2_O-10 U·ml^-1^ heparinized saline (∼0.4-0.5 v/v) and intravenously infused throughout the experiment to maintain hematocrit. Mice were euthanized by cervical dislocation immediately following the final arterial blood sample. Plasma and freeze-clamped tissue samples were stored at -20°C and - 80°C, respectively. Stable isotopes were purchased from Cambridge Isotope Laboratories Inc. (Tewksbury, MA).

### 2.5 Glucose derivatization and gas-chromatography-mass spectrometry (GC-MS) analysis

Approximately 40 μl of plasma acquired before starting the stable isotope infusions and at 90, 100, and 110 minutes following the [^13^C_3_]propionate bolus was used to prepare di-*O*-isopropylidene propionate, aldonitrile pentapropionate, and methyloxime pentapropionate derivatives of glucose, which were then analyzed via GC-MS as previously described [20, 38, 39]. Briefly, GC-MS protocols were performed using an Agilent 7890A gas chromatography system with an HP-5 ms capillary column (19091S-433, Agilent Technologies Inc., Santa Clara, CA) coupled to a 5975C mass spectrometer (Agilent Technologies Inc., Santa Clara, CA). The MS was run in scan mode for methyloxime (*m/z* 140-260), aldonitrile (*m/z* 100-500), and di-*O*-isopropylidene derivatives (*m/z* 301-314). Each derivative peak was integrated via a custom MATLAB function to determine mass isotopomer distributions (MIDs) for six fragment ions: methyloxime, *m/z* 145–149; aldonitrile, *m/z* 173–178, 259–266, 284–291, and 370–379; di-*O*- isopropylidene, *m/z* 301–314.

### 2.6 ^2^H/^13^C metabolic flux analysis

The *in vivo* metabolic flux analysis used in this study has been detailed previously [17, 38]. Briefly, a reaction network was generated using Isotopomer Network Compartmental Analysis (INCA) software [40]. This reaction network defined both carbon and hydrogen transitions for endogenous glucose production and associated oxidative metabolism reactions. The flux through each network reaction was determined relative to citrate synthase flux (V_CS_) by minimizing the sum of squared residuals between experimentally determined and simulated MIDs of the six fragment ions previously described [41, 42]. Flux estimations were repeated 50 times from random initial values. Goodness-of-fit was accepted according to a chi-square test (*p*=0.0*5*) with 34 degrees of freedom. Flux values for each mouse were an average of estimates obtained from samples at steady state (90, 100, and 110 minutes following the [^13^C_3_]propionate bolus) and normalized to liver weight.

### 2.7 Pyruvate tolerance tests

For pyruvate tolerance tests (PTTs), mice were fasted for five hours and injected intraperitoneally (i.p.) with pyruvate at a dose of 2 g·kg^-1^ lean weight. Blood glucose via tail cut sampling was measured at 0, 15, 30, 45, 60, 90, and 120 minutes with a Contour blood glucose meter (Ascensia Diabetes Care, Parsippany, NJ).

### 2.8 Circulating hormone analyses

Plasma insulin was quantified using a radioimmunoassay as previously performed [43] with plasma collected at the 120-minute time point (8 hours of fasting) from mice undergoing stable isotope infusions. Plasma glucagon was assessed in samples obtained at the 120-minute time point (8 hours of fasting) from mice undergoing the stable isotope infusions using the Mercodia Glucagon ELISA (10-1281-01, Winston Salem, NC, USA).

### 2.9 Liver lipid, metabolite, and enzyme analysis

For initial experiments wherein WT and KO mice were fed a high-fat diet (60% kcal from fat; Bio-Serv, F3282, Flemington, NJ) from 6 to 12 weeks of age, liver TAGs, diacylglycerides, and total cholesterol were quantified via chromatographic methods as previously described [44]. Liver samples from these mice were also sent to Metabolon® (Metabolon Inc., Research Triangle Park, NC, USA) for untargeted metabolomics as previously described [17, 45]. For mice included in the HF-SAAR diet studies, liver TAGs were determined with the Triglycerides-Liquid Reagent Set (Pointe Scientific Inc., Lincoln Park, MI) as previously outlined [46]. Liver samples from these mice were sent to Human Metabolome Technologies America Inc. (Boston, MA) for targeted, quantitative metabolomic analyses as previously detailed [20]. Liver glycogen was determined by an enzymatic assay as previously outlined [46, 47]. Aminotransferase activity was measured using Alanine Transaminase (ab105134) and Aspartate Aminotransferase (ab105135) Activity Assay Kits (Abcam, Cambridge, MA, USA). All liver samples used in analyses were obtained from mice fasted for 8 hours.

### 2.10 Liver histology

Liver tissue was formalin-fixed, paraffin-embedded, sectioned (5 μm) and stained with Masson’s Trichrome Blue by the Vanderbilt Translational Pathology Shared Resource to provide a measure of collagen. Whole slide digital images (20 X magnification) were completed using a Leica SCN400 Slide Scanner (Leica Microsystems, Buffalo Grove, IL, USA). Leica SlidePath Digital Image Hub software (Leica Microsystems, Buffalo Grove, IL, USA) was used for quantification protocols as previously described [17]. Briefly, pixels of interest were selected manually to create a color definition file for each stain. The parameters of this file were then applied to the entire tissue section. The measured stained area algorithm was used and the percentage of stained area was calculated as 100·(positive area/total tissue area). Two to three liver sections per mouse were quantified.

### 2.11 Immunoblotting

Liver tissue homogenates were prepared as previously detailed [20, 37, 39]. Liver proteins (15 μg) were separated via gel electrophoresis on a NuPAGE 4-12% Bis-Tris gel (Invitrogen, Carlsbad, CA, USA) and transferred to a PVDF membrane. The primary and secondary antibodies used are provided in Supplementary Table S2. Following incubation with antibodies, the PVDF membranes were treated with a chemiluminescent substrate (Thermo Fisher Scientific, Waltham, MA, USA) and images were obtained using a ChemiDoc™ Imaging system and Image Lab™ software (Bio-Rad, Hercules, CA, USA). A BLOT-FastStain (G-Bioscience, St. Louis, MO, USA) assay determined total protein, which was used as the loading control. Densitometry was completed using ImageJ software.

### 2.12 Mitochondrial isolation and high-resolution respirometry

Liver mitochondria were isolated from 8-hour fasted mice as previously described [48]. High-resolution respirometry experiments were performed using an Oroboros Oxygraph-2k (Oroboros Instruments, Innsbruck, Austria) at 37°C in duplicate and 2 ml of MiR05 (0.5 mM EGTA, 3 mM MgCl_2_-6H_2_O, 20 mM taurine, 10 mM KH_2_PO_4_, 20 mM HEPES, 1 g·liter^-1^ BSA, 60 mM potassium-lactobionate, 110 mM sucrose, pH 7.1, adjusted at 30°C). State 2 (CI: PMG) oxygen flux was tested with NADH-linked substrates, pyruvate (5 mM), glutamate (10 mM), and malate (2 mM). State 3 (CI: PMGD) oxygen consumption was assessed by a 5 mM ADP addition. State 3 (CI+CII: PMGDS) oxygen flux was quantified following the addition of succinate (10 mM). The respiratory control ratio (RCR) was determined by State 3 (CI: PMGD)/State 2 (CI: PMG). The succinate control ratio (SCR) was determined by State 3 (CI+CII: PMGDS)/State 3 (CI: PMGD).

### 2.13 Statistical analyses

GraphPad Prism software (GraphPad Software LLC, San Diego, CA) was used to complete statistical analyses. Analyses were performed using Student’s t-tests, two-way ANOVAs, and two-way repeated measures ANOVAs. If a significant interaction was detected for ANOVAs, a Sidak’s post hoc test was performed. Statistical differences were considered significant if p<0.05. All data are reported as mean ± SEM. Data points greater than two standard deviations from the mean were considered outliers and excluded from analysis.

## 3. Results

### 3.1 Loss of GNMT dysregulates methionine cycle homeostasis and promotes fatty liver

Initial studies fed WT and KO mice a high-fat diet for 6 weeks to model the early stages of MASLD (Fig. 1A). Loss of GNMT was confirmed by immunoblotting (Fig. 1B). Untargeted metabolomics showed that GNMT deletion dysregulated liver methionine cycle homeostasis (Fig. 1C and D). Glycine was comparable between genotypes (Fig. 1D). However, liver methionine and SAM were increased in GNMT KO mice (Fig. 1D). This was accompanied by a decline in liver SAH and adenosine (Fig. 1D). Loss of GNMT also perturbed methionine cycle-related metabolites. Liver choline was lower, betaine was higher, and dimethylglycine (DMG) was elevated in GNMT KO mice (Fig. 1D).

**Figure 1.**
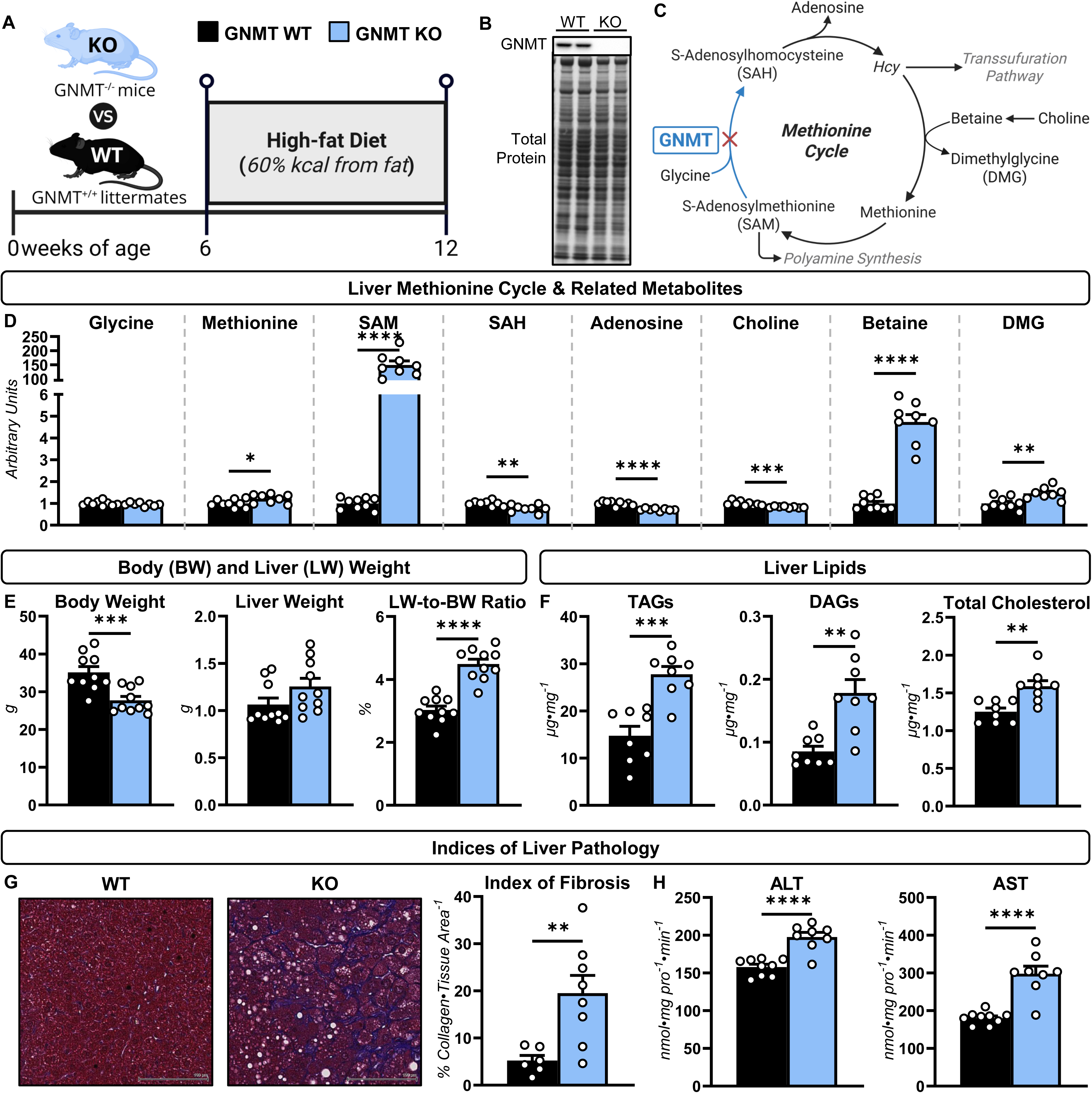
Loss of glycine N-methyltransferase (GNMT) promotes fatty liver in mice fed a high-fat diet. **A:** A schematic representation of the experimental design. GNMT knockout (KO) mice and wild type (WT) littermates were fed a high-fat diet for 6 weeks. **B:** A representative immunoblot of GNMT in livers of WT and KO mice. **C:** A schematic representation of the methionine cycle with select metabolites and enzymes. **D:** Liver glycine, methionine, s-adenosylmethionine (SAM), s-adenosylhomocysteine (SAH), adenosine, choline, betaine, and dimethylglycine (DMG) in 12-week-old mice (arbitrary units; n = 8-9 per genotype). **E:** Body weight (g), liver weight (g), and liver weight (LW)-to-body weight ratio (%) in 12-week-old WT and KO mice (n = 10 per genotype). **F:** Liver triacylglycerides (TAGs), diacylglycerides (DAGs), and total cholesterol (µg·mg^-1^) in 12-week-old WT and KO mice (n = 8 per genotype). **G:** Percentage of collagen area per tissue area as determined by Masson’s trichrome blue stain and representative images (×20 magnification) for WT and KO mice (n = 6-8 per genotype). **H:** Liver alanine aminotransferase (ALT) and aspartate aminotransferase (AST) activity (nmol·mg^-1^ pro·min^-1^; n = 8-9 per genotype). Data are mean ± SEM. Statistical differences were determined by a Student’s t test and accepted as significant if p<0.05. *p<0.05, **p<0.01, ***p<0.001, and ****p<0.0001.

The perturbations in methionine homeostasis were associated with features of systemic and liver-specific pathophysiology (Fig. 1E-H). Body weight was lower in mice lacking GNMT (Fig. 1E), which was due to lower adiposity (Fig. S1A). Liver weight did not change; however, the liver-to-body weight ratio was higher in GNMT KO mice (Fig. 1E). Loss of GNMT increased liver TAGs (∼1.9-fold), DAGs (∼2.1-fold), and total cholesterol (∼1.3-fold) (Fig. 1F). Collagen as a percentage of liver tissue area was significantly higher in GNMT KO mice compared to WT littermates (Fig. 1G). The enzymatic activities of alanine (ALT) and aspartate (AST) aminotransferase were elevated in livers of mice lacking GNMT (Fig. 1H). Together, these results demonstrate that the loss of GNMT promotes liver steatosis, fibrosis, and injury in mice fed a high-fat diet.

### 3.2 GNMT KO mice exhibit impaired endogenous glucose production

The current study tested the hypothesis that loss of GNMT-mediated transmethylation inhibits liver glucose formation in mice fed a high-fat diet. Following 6 weeks of high-fat feeding, ^2^H- and ^13^C-isotope infusions in conscious, unrestrained mice were completed to quantify glucose fluxes *in vivo* (Fig. 2A). Plasma insulin was lower (Fig. 2B), glucagon trended higher (Fig. 2C), and steady-state glucose concentrations were similar (Fig. 2D) in KO mice compared to WT mice. Liver glucose precursors were perturbed by loss of GNMT. Glycogen was decreased (Fig. 2E), glycerol increased (Fig. 2F), and phosphoenolpyruvate (PEP; Fig. 2G) trended lower in livers of KO mice. These changes in static metabolites were consistent with glucose fluxes. Endogenous glucose production (V_EGP_) decreased in KO mice (Fig. 2H). Glycogenolysis, flux from glycogen to glucose (V_PYGL_), was inhibited in mice lacking GNMT (Fig. 2I). Gluconeogenesis from glycerol (V_GK_) was increased; however, lower gluconeogenic flux from PEP (V_Enol_) resulted in impaired total gluconeogenesis (V_Aldo_; Fig. 2J-L).

**Figure 2.**
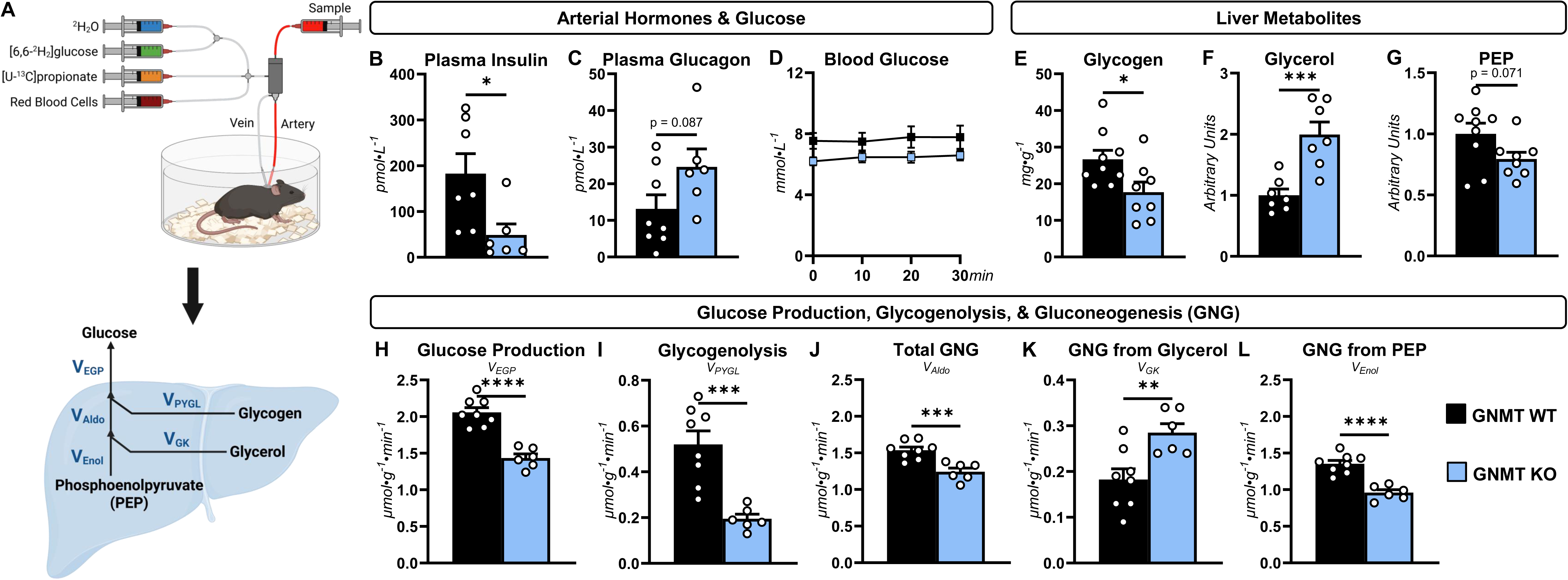
Endogenous glucose production is impaired in glycine N-methyltransferase (GNMT) knockout (KO) mice. **A:** GNMT KO mice and wild type (WT) littermates fed a high-fat diet for 6 weeks underwent ^2^H- and ^13^C-isotope infusions to quantify endogenous glucose fluxes. **B:** Plasma insulin (pmol·L^-1^; n = 6-7 per genotype). **C:** Plasma glucagon (pmol·L^-1^; n = 6-8 per genotype). **D:** A time course of arterial blood glucose during acquisition of plasma for ^2^H/^13^C metabolic flux analysis (mmol·L^-1^; n = 6-8 per genotype). **E:** Liver glycogen (mg·g^-1^; n = 8-9 per genotype). **F:** Liver glycerol (arbitrary units; n = 7 per genotype). **G:** Liver phosphoenolpyruvate (PEP; arbitrary units; n = 8-9 per genotype). Model-estimated, absolute nutrient fluxes (μmol·g^-1^·min^-1^) in WT and GNMT KO mice (n = 6-8 per genotype) for **H:** endogenous glucose production (V_EGP_), **I:** glycogenolysis (V_PYGL_), **J:** total gluconeogenesis (GNG; V_Aldo_), **K:** GNG from glycerol (V_GK_), and **L:** GNG from PEP (V_Enol_). Data are mean ± SEM. Statistical differences were determined by a Student’s t test and accepted as significant if p<0.05 for data in panels B-C and E-L. Statistical differences (p<0.05) were tested by a two-way repeated measures ANOVA in panel D. *p<0.05, **p<0.01, ***p<0.001, and ****p<0.0001.

### 3.3 Lower partitioning of TCA cycle intermediates to glucose in mice lacking GNMT

The generation of PEP for gluconeogenic flux is linked to the entry and exit of nutrients into and from the TCA cycle (Fig. 3A). Pyruvate cycling, the flux of PEP to pyruvate (V_PK+ME_), was comparable between genotypes (Fig. 3B). TCA cycle cataplerosis (V_PCK_; flux from oxaloacetate to PEP) was lower in KO mice (Fig. 3C). Anaplerotic fluxes, V_PC_ (flux from pyruvate to oxaloacetate; Fig. 3D), V_LDH_ (flux of unlabeled lactate to pyruvate; Fig. 3E), and V_PCC_ (flux from propionyl-CoA to succinyl-CoA; Fig. 3F) were impaired in mice lacking GNMT. In contrast to cataplerotic and anaplerotic fluxes, TCA cycle fluxes (V_CS_ and V_SDH_) were similar between WT and KO mice (Fig. 3G and H). Notably, multiple TCA cycle metabolites were higher in livers of KO mice (Fig. 3I). These metabolites included citrate, aconitate, succinylcarnitine (surrogate for succinyl-CoA), and propionylcarnitine (surrogate for propionyl-CoA) (Fig. 3I). These results indicate that loss of GNMT inhibits TCA cycle nutrient cataplerosis for use in gluconeogenesis.

**Figure 3.**
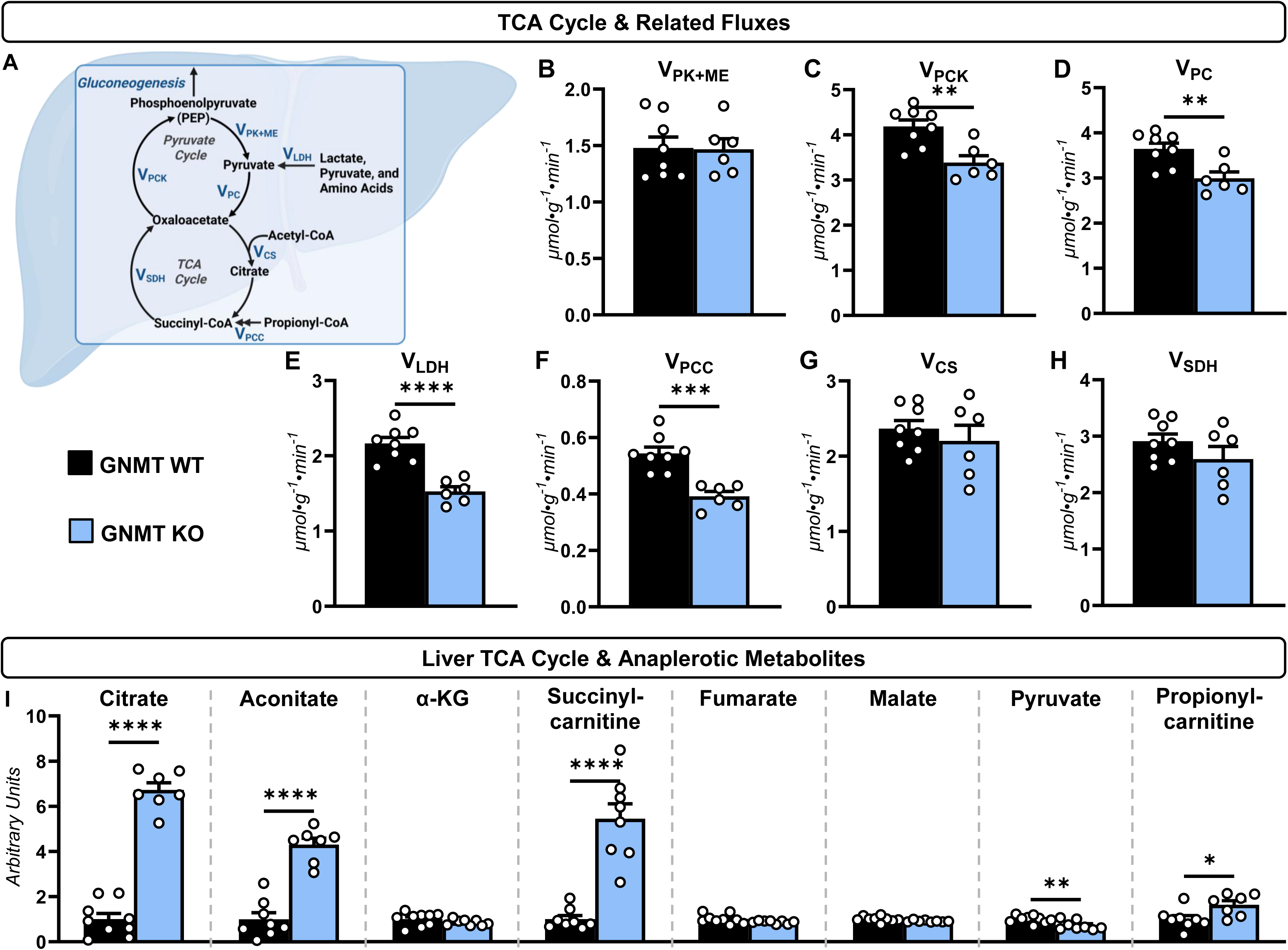
Cataplerotic flux to glucose is impaired, but tricarboxylic acid cycle flux is preserved in glycine N-methyltransferase (GNMT) knockout (KO) mice fed a high-fat diet. **A:** A schematic representation of TCA cycle and related fluxes in 12-week-old wild type (WT) and GNMT KO mice fed a high-fat diet for 6 weeks. Nutrient fluxes (μmol·g^-1^·min^-1^; n = 6-8 per genotype) quantified were **B:** pyruvate cycling (V_PK+ME_), **C:** tricarboxylic acid cycle cataplerosis to phosphoenolpyruvate (V_PCK_), **D:** anaplerosis from pyruvate (V_PC_), **E:** flux from unlabeled, anaplerotic sources to pyruvate (V_LDH_), **F:** anaplerosis from propionyl-CoA (V_PCC_), **G:** flux from oxaloacetate and acetyl-CoA to citrate (V_CS_), and **H:** flux from succinyl-CoA to oxaloacetate (V_SDH_). **I:** Liver citrate, aconitate, α-ketoglutarate (α-KG), succinylcarnitine, fumarate, malate, pyruvate, and propionylcarnitine (arbitrary units; n = 7-9 per genotype). Data are mean ± SEM. Statistical differences were determined by a Student’s t test and accepted as significant if p<0.05. *p<0.05, **p<0.01, ***p<0.001, and ****p<0.0001.

### 3.4 Indices of lipid synthesis, desaturation, and elongation are increased in KO mice

TCA cycle intermediates may be used in biosynthetic processes beyond glucose production. For example, citrate can be used for the production of acetyl-CoA, and subsequently, fatty acids via *de novo* lipogenesis (Fig. 4A). Given this, we assessed molecular regulators of lipogenesis in livers of WT and KO mice (Fig. 4B). Liver ATP-citrate lyase (ACLY) and fatty acid synthase protein levels were comparable between genotypes (Fig. 4B). Liver acetyl-CoA carboxylase (ACC) and stearoyl-CoA desaturase 1 (SCD1) were elevated in KO mice (Fig. 4B). Consistent with the increased ACC protein, liver malonyl-carnitine (surrogate for malonyl-CoA) was ∼6-fold higher in KO mice compared to WT littermates (Fig. 4C). In agreement with the higher SCD1 expression, loss of GNMT led to elevated desaturation of fatty acids comprising liver TAGs. Specifically, liver C16:1 and C18:1 concentrations and desaturation indexes (C16:1-to-C16:0 and C18:1-to-C18:0 ratios) were increased in KO mice (Fig. 4D and E). The elongation index [(C18:0 + C18:1)/C16:0] of fatty acids comprising liver TAGs was also higher in mice lacking GNMT (Fig. 4F). The results suggest that livers of GNMT KO mice exhibit increased fatty acid synthesis, desaturation, and elongation. Furthermore, loss of GNMT may promote fatty liver by redirecting the fate of TCA cycle intermediates (and acetyl-CoA) away from supporting gluconeogenesis and towards lipid deposition.

**Figure 4.**
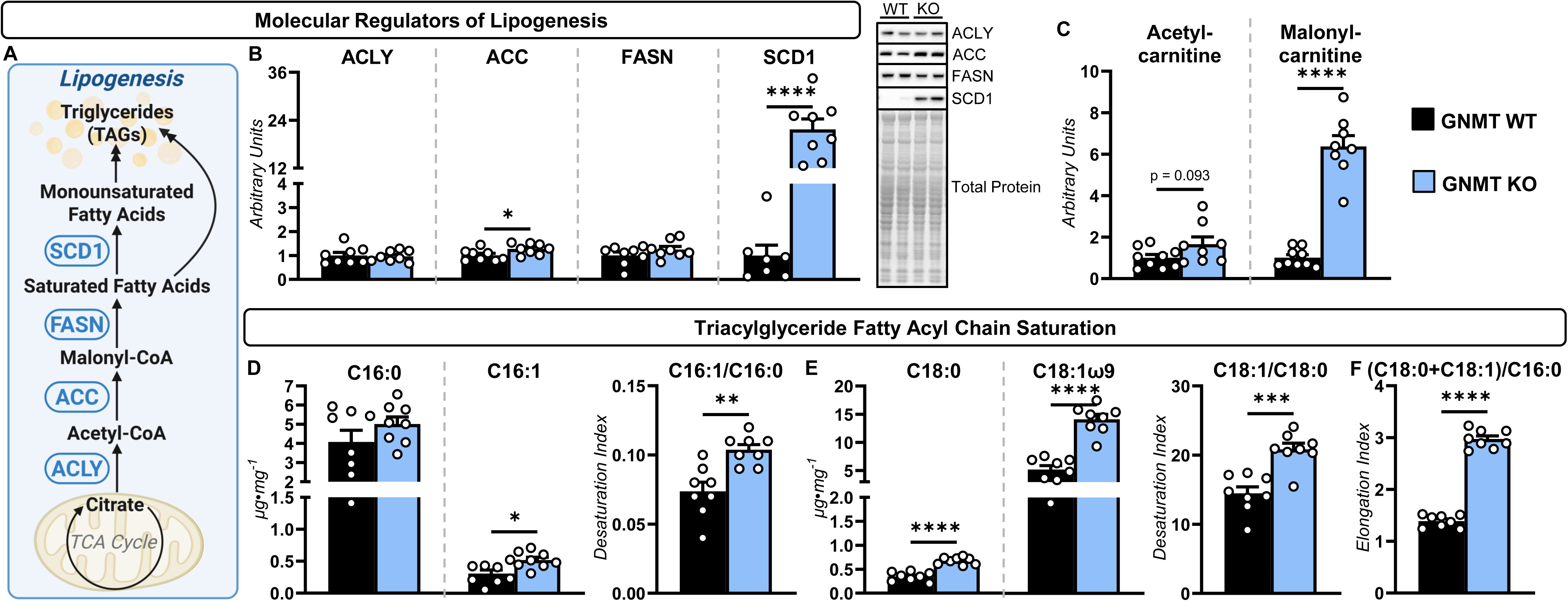
Indices of lipid synthesis, desaturation, and elongation are increased in glycine N-methyltransferase (GNMT) knockout (KO) mice. Twelve-week-old wild type (WT) and GNMT KO mice were fed a high-fat diet for 6 weeks prior to analyses. **A:** A schematic representation of *de novo* lipogenesis. **B:** Liver ATP-citrate lyase (ACLY), acetyl-CoA carboxylase (ACC), fatty acid synthase (FASN), and stearoyl-CoA desaturase 1 (SCD1) as determined by immunoblotting and representative immunoblots (arbitrary units; n = 7-8 per genotype). **C:** Liver acetylcarnitine and malonylcarnitine (arbitrary units; n = 8-9 per genotype). **D:** C16:0 and C16:1 fatty acids in liver TAGs (µg·mg^-1^) and the C16:1-to-C16:0 ratio (n = 7-9 per genotype). **E:** C18:0 and C18:1 fatty acids in liver TAGs (µg·mg^-1^) and the C18:1-to-C18:0 ratio (n = 8 per genotype). **F:** Elongation index [(C18:0+C18:1)/C16:0] of fatty acyl chains in liver TAGs (n = 8 per genotype). Data are mean ± SEM. Statistical differences were determined by a Student’s t test and accepted as significant if p<0.05. *p<0.05, **p<0.01, ***p<0.001, and ****p<0.0001.

### 3.5 Increased SAM is linked to metabolic remodeling in GNMT KO mice

Beyond its role as a methyl donor, SAM regulates and is used in multiple metabolic processes. This includes promoting transsulfuration via allosteric activation of cystathionine-β-synthase (CBS) [49, 50]. Consistent with allosteric activation of transsulfuration, the protein levels of transsulfuration enzymes, CBS and cystathionine γ-lyase (CSE), were similar in the livers of WT and KO mice (Fig. 5A and B). However, livers of KO mice exhibited decreased serine, increased cystathionine, and higher cysteine (Fig. 5A and C). Notably, cysteine is a precursor for glutathione synthesis (Fig. 5A). Both reduced (GSH) and oxidized (GSSG) glutathione were elevated in livers of KO mice (Fig. 5A and C). This increase in glutathione was accompanied by higher glutathione synthetase (GSS) expression (Fig. 5A and D). The connection between elevated SAM and glutathione is notable because recent work has demonstrated that this antioxidant positively regulates liver lipid synthesis by promoting the expression and stability of *de novo* lipogenesis enzymes [51, 52]. In speculation, the increased SAM in KO mice may stimulate *de novo* lipogenesis by elevating glutathione levels.

**Figure 5.**
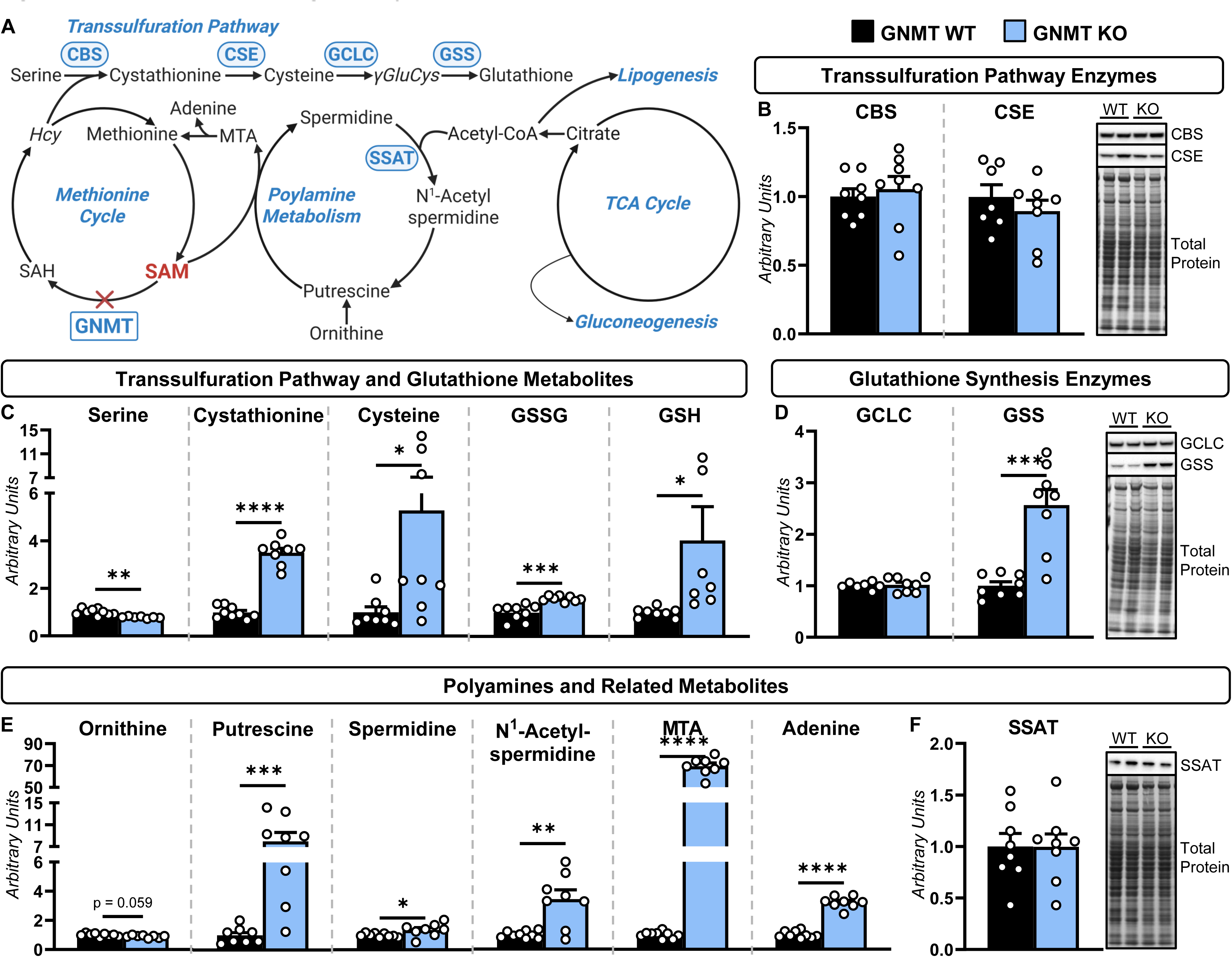
Altered SAM-consuming pathways in livers of glycine N-methyltransferase (GNMT) knockout (KO) mice. Twelve-week-old wild type (WT) and GNMT KO mice were fed a high-fat diet for 6 weeks prior to analyses. **A:** A schematic of the enzymes, metabolites and pathways connected to s-adenosylmethionine (SAM) availability. **B:** Liver cystathionine-β-synthase (CBS) and cystathionine γ-lyase (CSE) as determined by immunoblotting and representative immunoblots (arbitrary units; n = 7-8 per genotype). **C:** Liver serine, cystathionine, cysteine, oxidized glutathione (GSSG), and reduced glutathione (GSH) (arbitrary units; n = 7-9 per genotype). **D:** Liver glutamate-cysteine ligase catalytic subunit (GCLC) and glutathione synthetase (GSS) as determined by immunoblotting and representative immunoblots (arbitrary units; n = 7-8 per genotype). **E:** Liver ornithine, putrescine, spermidine, N^1^-acetylspermidine, 5’-methylthioadenosine (MTA), and adenine (arbitrary units; n = 8-9 per genotype). **F:** Liver spermidine/spermine N^1^-acetyltransferase (SSAT) as determined by immunoblotting and representative immunoblots (arbitrary units; n = 8 per genotype). Data are mean ± SEM. Statistical differences were determined by a Student’s t test and accepted as significant if p<0.05. *p<0.05, **p<0.01, ***p<0.001, and ****p<0.0001. Homocysteine, Hcy; s-adenosylhomocysteine, SAH; gamma-glutamylcysteine, γGluCys. Italicized metabolites were not determined.

The results of untargeted metabolomics also suggest that excess SAM is directed towards polyamine synthesis (Fig. 5A and E). Liver putrescine, spermidine, and 5-methylthioadenosine (MTA) were increased and ornithine was reduced in KO mice (Fig. 5E). Liver adenine, a product of MTA catabolism, was elevated in mice lacking GNMT (Fig. 5E). Polyamine concentrations are also regulated by a catabolic pathway involving spermidine acetylation, which facilitates the resynthesis of putrescine and may assist in disposing of excess SAM [53]. N^1^-acetylspermidine was higher in KO mice (Fig. 5E). However, spermidine/spermine N-acetyltransferase (SSAT) protein was unchanged between genotypes (Fig. 5A and F). Together, these data suggest that the accumulation of SAM in GNMT KO mice may promote a futile cycle of polyamine synthesis and catabolism. As a result, the persistent elevation in liver SAM may “pull” acetyl-CoA into polyamine acetylation reactions and away from oxidation in the TCA cycle. The increased polyamines in GNMT KO mice are of particular interest for two reasons. First, enhancing polyamine acetylation via SSAT overexpression increases energy expenditure and lowers adiposity [54]. We show that the lower adiposity in GNMT KO mice is accompanied by elevated energy expenditure (Fig. S1). Second, and more importantly, polyamines promote tumor initiation and progression [55].

### 3.6 Dietary SAAR prevents increased SAM availability and liver pathology in KO mice

Prior work has determined that GNMT KO mice develop HCC by 32 weeks of age [21]. To test if this is owing to the accumulation of SAM, WT and KO mice were fed a HF-CTRL or HF-SAAR diet from 6 to 32 weeks of age to prevent the accumulation of SAM (Fig. 6A and B). Body weight gain was less pronounced in KO mice compared to WT littermates fed a HF-CTRL diet (Fig. 6C). Dietary SAAR attenuated the gain in body weight in both genotypes (Fig. 6C). On a HF-CTRL diet, KO mice showed reduced adiposity compared to WT mice (Fig. 6D). A main effect for genotype and diet indicated that both loss of GNMT and dietary SAAR lowered lean mass (Fig. 6D). In addition to systemic adaptations, SAAR elicited liver-specific responses in mice (Fig. 6E-L). Specifically, the HF-SAAR diet increased liver glycine levels (Fig. 6F). SAAR prevented the increase in methionine, SAM, and SAH in KO mice (Fig. 6F). SAAR decreased liver weight in both genotypes and prevented the increase in the liver weight-to-body weight ratio in KO mice (Fig. 6G and H). Hydroxyproline, a marker of liver fibrosis, was higher in livers of KO mice fed a HF-CTRL diet compared to WT mice fed a HF-CTRL diet; however, the rise in KO mice was prevented by the HF-SAAR diet (Fig. 6I). In contrast to 12-week-old mice fed a HF-CTRL diet, 32-week-old KO mice had lower liver TAGs relative to WT mice receiving the HF-CTRL diet (Fig. 6J). The HF-SAAR diet decreased liver TAGs in both genotypes (Fig. 6J). The HCC marker, minichromosome maintenance complex component 2 (MCM2), was elevated in KO mice versus WT mice fed the HF-CTRL diet (Fig. 6K). This was accompanied by tumor nodules in livers of KO mice receiving the HF-CTRL diet (Fig. 6L). Dietary SAAR abolished the rise in liver MCM2 protein and appearance of tumor nodules in KO mice (Fig. 6K and L). Together, these results suggest that the SAAR diet attenuated the increase in SAM availability and associated liver pathology in KO mice.

**Figure 6.**
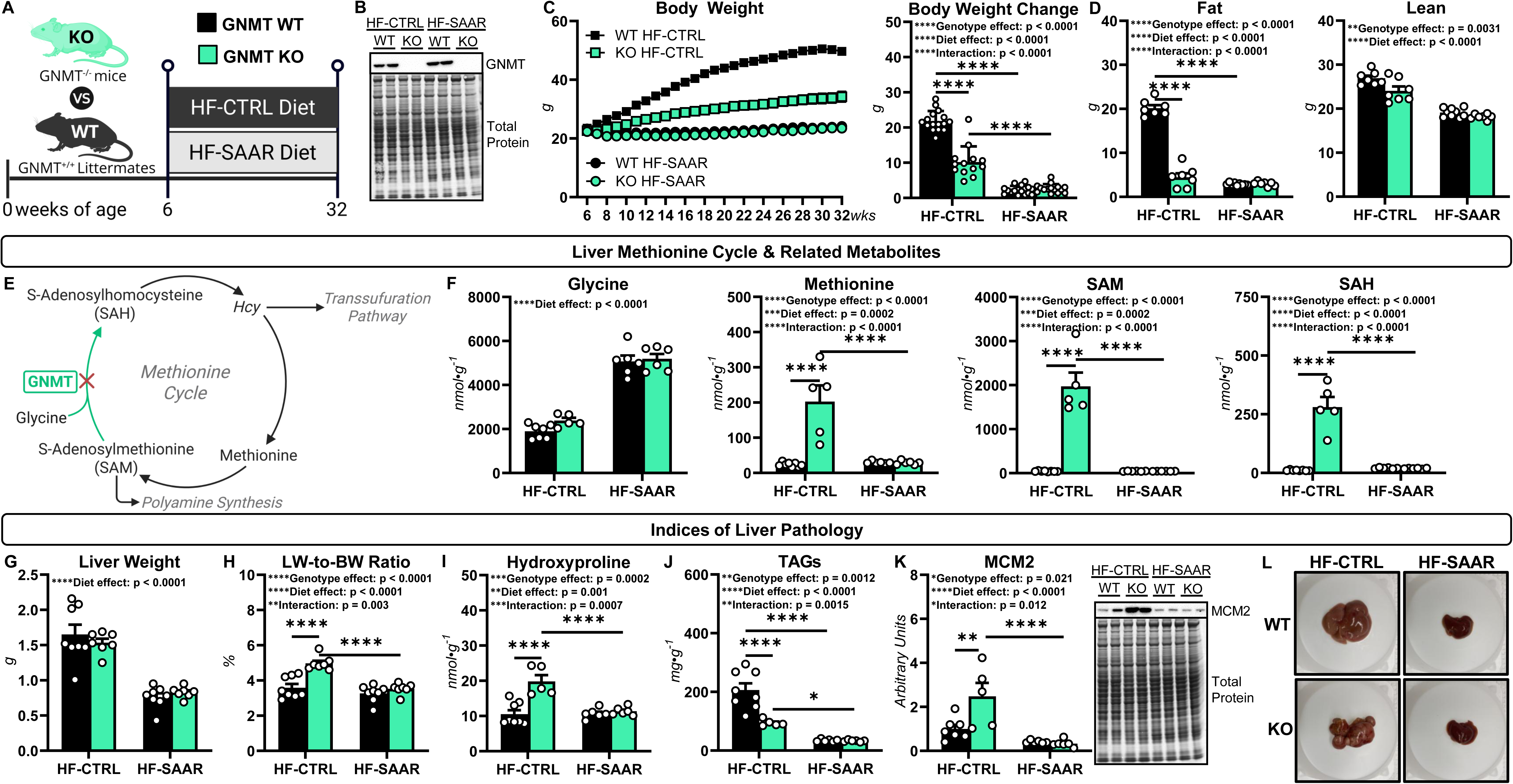
Dietary sulfur amino acid restriction (SAAR) prevents increased liver S-adenosylmethionine (SAM) and pathology in glycine N-methyltransferase (GNMT) knockout (KO) mice. **A:** A schematic representation of the experimental design. GNMT knockout (KO) mice and wild type (WT) littermates were fed a high-fat control (HF-CTRL) or HF-SAAR diet for 26 weeks starting at 6 weeks of age. **B:** A representative immunoblot of GNMT in livers of WT and KO mice. **C:** A body weight time course and change in body weight over the 26-week dietary protocol (g; n = 13-16 per group). **D:** Fat and lean mass in WT and KO mice after the 26-week dietary protocol (g; n = 7-9 per group). **E:** A schematic representation of the methionine cycle with select metabolites and enzymes. **F:** Liver glycine, methionine, SAM, and s-adenosylhomocysteine (SAH) following the 26-week HF-CTRL and HF-SAAR feeding (nmol·g^-1^; n = 5-8 per group). **G:** Liver weight (g; n = 7-8 per group). **H:** Liver weight (LW)-to-body weight (BW) ratio (%; n = 7-8 per group). **I:** Liver hydroxyproline (nmol·g^-1^; n = 5-8 per group). **J:** Liver triacylglycerides (TAGs; mg·g^-1^; n = 5-8 per group). **K:** Liver minichromosome maintenance complex component 2 (MCM2) as determined by immunoblotting and representative immunoblots (arbitrary units; n = 5-8 per group). **L:** Representative image of livers from WT and KO mice. Data are mean ± SEM. Statistical differences (p<0.05) were determined by a two-way ANOVA followed by Sidak’s post hoc tests. Significant main and/or interaction effects are presented within each panel. *p<0.05, **p<0.01, and ****p<0.0001. Homocysteine, Hcy; s-adenosylhomocysteine. Italicized metabolites were not determined.

### 3.7 SAAR limits metabolic perturbations in livers of GNMT KO mice

Next, we tested the efficacy of dietary SAAR to mitigate metabolic perturbations in GNMT KO mice. Liver glycogen was higher in KO mice compared to WT mice fed a HF-CTRL diet (Fig. 7A). A SAAR diet prevented differences in liver glycogen between genotypes (Fig. 7A). Given these changes in glycogen, we assessed intermediates of glycogen synthesis and breakdown. A main effect of diet indicated that dietary SAAR increased glucose-6-phosphate (G6P) and glucose-1-phosphate (G1P) (Fig. 7B). Notably, SAAR lowered UDP-glucose, the immediate precursor of glycogen, in KO mice (Fig. 7B).

**Figure 7.**
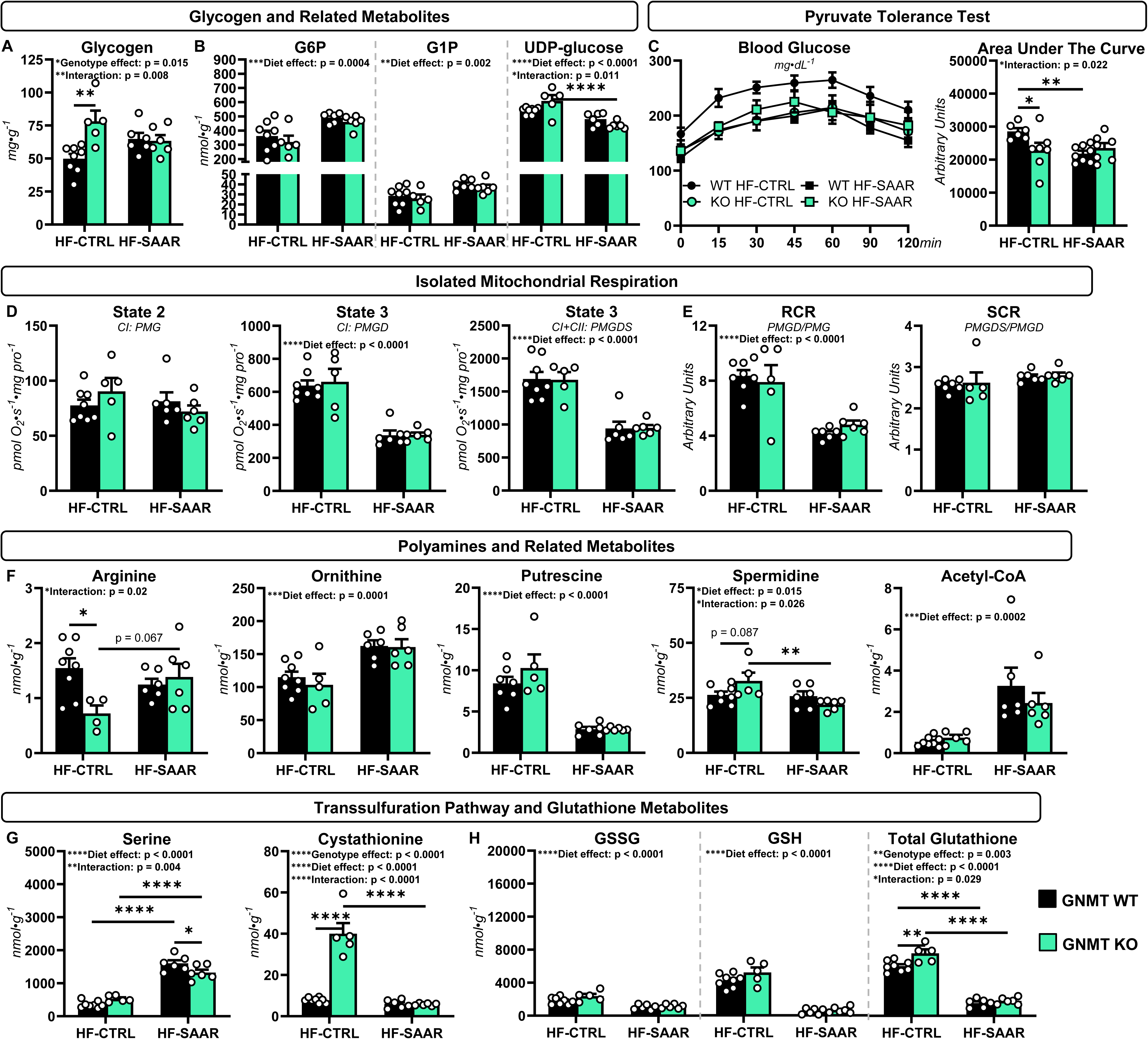
Long-term dietary sulfur amino acid restriction (SAAR) mitigates liver metabolic remodeling in glycine N-methyltransferase (GNMT) knockout (KO) mice. GNMT knockout (KO) mice and wild type (WT) littermates were fed a high-fat control (HF-CTRL) or HF-SAAR diet for 26 weeks starting at 6 weeks of age. **A:** Liver glycogen (mg·g^-1^; n = 5-8 per group) **B:** Liver glucose-6-phosphate (G6P), glucose-1-phosphate (G1P), and UDP-glucose (nmol·g^-1^; n = 5-8 per group). **C:** Glucose excursion and area under the curve during a pyruvate tolerance test (n = 6-8 per group). **D:** Isolated liver mitochondria oxygen consumption rates normalized to mitochondrial protein (n = 5-8 per group; pmol O_2_·s^-1^·mg^-1^). **E:** Respiratory control ratio (RCR; n = 5-8) and succinate control ratio (SCR; n = 5-7 per group). **F:** Liver arginine, ornithine, putrescine, spermidine, and acetyl-CoA (nmol·g^-1^; n = 5-8 per group). **G:** Liver serine, cystathionine, oxidized glutathione (GSSG), reduced glutathione (GSH), and total glutathione (GSSG+GSH) (nmol·g^-1^; n = 5-8 per group). Data are mean ± SEM. Statistical differences (p<0.05) were determined by a two-way ANOVA followed by Sidak’s post hoc tests. Significant main and/or interaction effects are presented within each panel. *p<0.05, **p<0.01, ***p<0.001, and ****p<0.0001.

PTTs were completed to provide an index of gluconeogenesis (Fig. 7C). On a HF-CTRL diet, GNMT KO mice had reduced glucose excursion during PTTs compared to WT mice (Fig. 7C). Glucose excursion in response to pyruvate administration was also lower in WT mice fed a HF-SAAR diet compared to WT mice fed the HF-CTRL diet (Fig. 7C). Liver mitochondrial function was assessed via isolated mitochondrial respirometry (Fig. 7D and E). State 2 respiration supported by NADH-linked substrates was comparable between all groups (Fig. 7D). Dietary SAAR lowered State 3 respiration supported by NADH-linked substrates as well as State 3 respiration supported by combined NADH- and FADH_2_-linked substrates (Fig. 7D). The respiratory control ratio was comparable between genotypes within each diet; however, dietary SAAR reduced RCR in both genotypes (Fig. 7E). The SCR was comparable between all groups (Fig. 7E).

Targeted metabolomics was completed to evaluate metabolite levels within polyamine turnover and transsulfuration pathways. Dietary SAAR prevented the decline in liver arginine, a polyamine precursor, in KO mice (Fig. 7F). The HF-SAAR diet increased ornithine and decreased putrescine in both genotypes (Fig. 7F). KO mice fed a HF-SAAR diet also showed lower liver spermidine compared to KO mice receiving the HF-CTRL diet (Fig. 7F). This was accompanied by increased liver acetyl-CoA in mice fed the HF-SAAR diet (Fig. 7F). Together, these results suggest that the SAAR diet may attenuate increased polyamine synthesis and catabolism in KO mice. Assessment of transsulfuration pathway metabolites showed liver serine was comparable between genotypes on the HF-CTRL diet (Fig. 7G). The HF-SAAR diet elevated liver serine; however, this increase was blunted in KO mice (Fig. 7G). KO mice fed the HF-CTRL diet displayed higher liver cystathionine compared to WT controls (Fig. 7G). Dietary SAAR attenuated the rise in liver cystathionine (Fig. 7G). Total liver glutathione was elevated in KO mice relative to WT mice on a HF-CTRL diet (Fig. 7H). The HF-SAAR diet decreased total glutathione in both genotypes (Fig. 7H). These results suggest that SAAR limits transsulfuration and glutathione synthesis in livers of mice lacking GNMT.

### 3.8 SAAR mitigates fatty liver and molecular regulators of lipogenesis in KO mice

To determine if a HF-SAAR diet prevented the fatty liver in GNMT KO mice prior to the appearance of HCC, WT and KO mice were fed a HF-CTRL or HF-SAAR diet for 6 weeks starting at 6 weeks of age (Fig. 8A and B). Consistent with our previous results, loss of GNMT reduced body weight and adiposity compared to WT mice fed a HF-CTRL diet (Fig. 8C and D). Dietary SAAR decreased body weight in both genotypes (Fig. 8C) and fat mass in WT mice (Fig. 8D). Lean mass was lower in mice fed the HF-SAAR diet as indicated by a main effect for diet (Fig. 8E). On the HF-CTRL diet, liver TAGs were elevated in KO compared to WT mice (Fig. 8F). Notably, the HF-SAAR diet lowered liver TAGs in both genotypes (Fig. 8F). This was accompanied by SAAR preventing the increase in liver ACC, FASN, and SCD1 in GNMT KO mice (Fig. 8G). These results indicate that dietary SAAR may prevent the initiation and/or progression to HCC in response to GNMT deletion by blocking the liver steatosis that precedes HCC.

**Figure 8.**
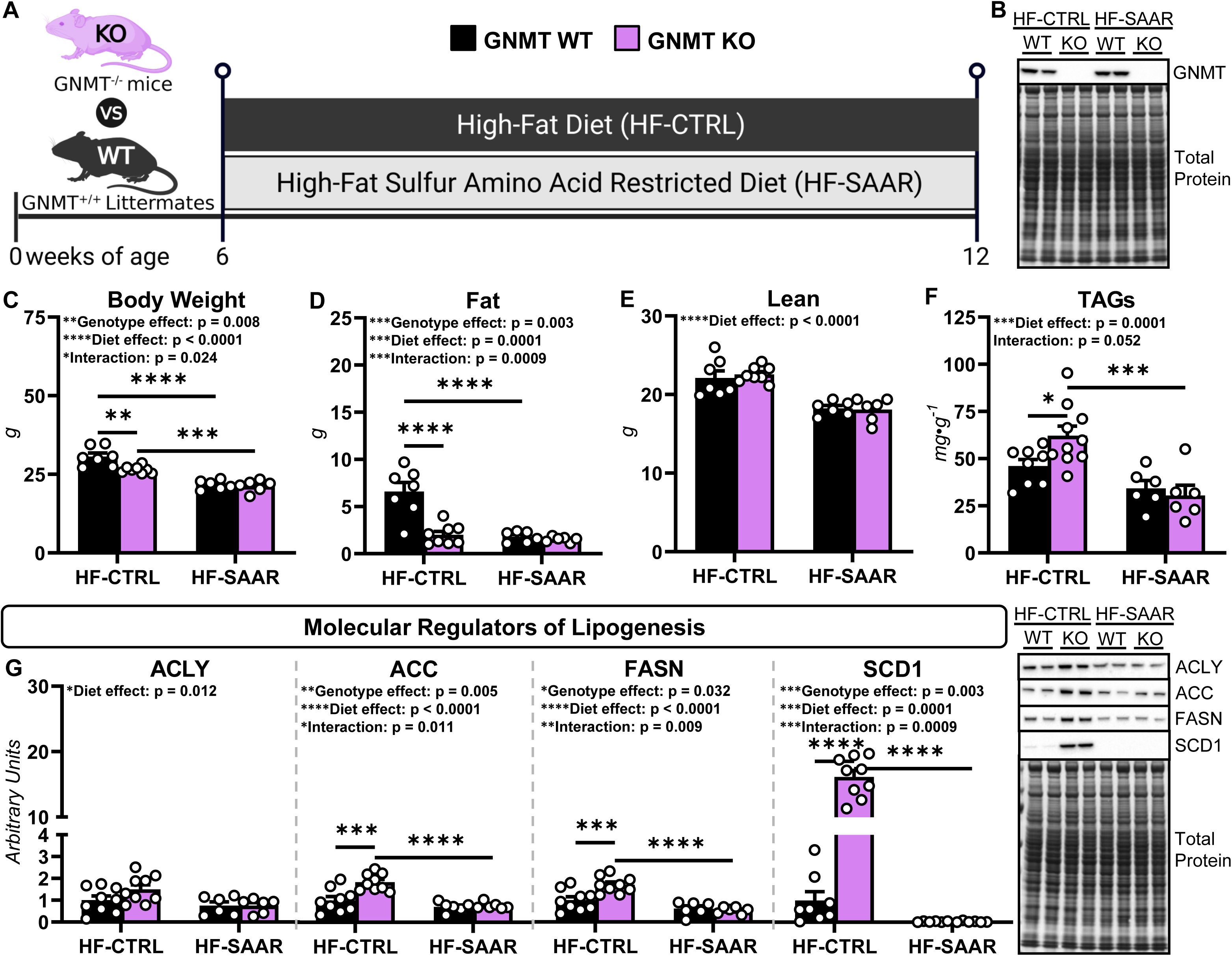
Short-term dietary sulfur amino acid restriction (SAAR) mitigates fatty liver in glycine N-methyltransferase (GNMT) knockout (KO) mice. **A:** A schematic representation of the experimental design. GNMT knockout (KO) mice and wild type (WT) littermates were fed a high-fat control (HF-CTRL) or HF-SAAR diet for 6 weeks starting at 6 weeks of age. **B:** A representative immunoblot of GNMT in livers of WT and KO mice. **C:** Body weight (g; n = 6-8 per group). **D:** Fat mass (g; n = 6-8 per group). **E:** Lean mass (g; n = 6-8 per group). **F:** Liver triacylglycerides (TAGs; mg·g^-1^; n = 6-10 per group). **G:** Liver ATP-citrate lyase (ACLY), acetyl-CoA carboxylase (ACC), fatty acid synthase (FASN), and stearoyl-CoA desaturase 1 (SCD1) as determined by immunoblotting and representative immunoblots (arbitrary units; n = 6-9 per group). Data are mean ± SEM. Statistical differences (p<0.05) were determined by a two-way ANOVA followed by Sidak’s post hoc tests. Significant main and/or interaction effects are presented within each panel. *p<0.05, **p<0.01, ***p<0.001, and ****p<0.0001.

## 4. Discussion

Prior work using both human samples and mouse models suggests that GNMT insufficiency promotes MASLD and HCC [21, 22, 27, 28]. Our previous work identified that liver glucose production via glycogenolysis and TCA cycle-supported gluconeogenesis is impaired in both 12-week-old GNMT-null mice with fatty liver and 44-week-old GNMT-null mice with HCC [11, 17, 20]. Importantly, a decline in liver glucose production has been hypothesized to support the development and maintenance of HCC by sparing nutrients and/or energy resources for tumorigenic processes [17, 56–58]. Here, we extend our prior studies performed in lean, low-fat-fed mice by quantifying liver glucose production and associated TCA cycle fluxes in mice with diet-induced obesity. Moreover, our work tested the extent to which elevated SAM mediates the metabolic phenotypes and MASLD-related HCC in GNMT KO mice. The key findings of this study are i) loss of GNMT in high-fat fed mice inhibits gluconeogenesis from TCA cycle intermediates, but does not diminish TCA cycle flux; ii) precursors are diverted away from gluconeogenesis to other biosynthetic processes in response to GNMT insufficiency as indicated by metabolite concentrations in pathways that use both SAM and TCA cycle intermediates; and iii) the appearance of fatty liver and HCC in high-fat-fed GNMT KO mice is dependent on increased SAM.

### 4.1 Glucose production and TCA cycle flux in the livers of GNMT KO mice

Here we show that 12-week-old GNMT KO mice fed a high-fat diet for 6 weeks displayed decreased endogenous glucose production (V_EGP_). This is consistent with our prior work in low-fat-fed GNMT-null mice prior to (12 weeks of age) and in the presence of HCC (44 weeks of age) [11, 17, 20]. The lower glucose formation in this study was, in part, due to a diminished contribution of glycogen to glucose synthesis (V*_PYGL_*). In agreement, impaired glycogenolysis was previously observed in lean GNMT-deficient mice [11, 17, 20]. Interestingly, these prior reports show that the decreased glycogenolysis is frequently accompanied by increased glycogen levels [17, 19, 20, 59], indicating that loss of GNMT in low-fat-fed mice inhibits the mobilization of glycogen for glucose production. In contrast, liver glycogen was lower in 12-week-old, high-fat-fed GNMT KO mice. This suggests that the decline in glycogenolysis in high-fat-fed KO mice, prior to the appearance of HCC, was not owing to impaired glycogen breakdown, but the result of limited glycogen availability. This distinction is important because it suggests two different mechanisms are preventing glycogen-derived glucose production depending on diet: substrate availability in high-fat-fed mice versus impaired mobilization in low-fat-fed mice. The lower glycogen in 12-week-old, high-fat-fed KO mice may be due to hormonal responses. Circulating insulin was lower and glucagon trended higher in KO mice, which could limit glycogen accumulation. Nevertheless, the work presented here indicates that loss of GNMT leads to decreased glycogenolysis in mice fed both a low- and high-fat diet prior to the appearance of HCC. However, the underlying mechanisms driving the reduced flux differ by dietary condition. It should be noted that this study also fed GNMT KO mice a high-fat diet from 6-to 32-weeks of age, when HCC is observed. Under these conditions, liver glycogen was higher in KO mice. This is consistent with prior reports demonstrating that abnormal hepatic glycogen storage is linked to increased risk of HCC [60, 61]. Further work is needed to determine whether the time-dependent switch from lower to higher liver glycogen in high-fat-fed KO mice plays a causal role in the development of HCC.

Concurrent with the reduced glycogenolysis, 12-week-old, high-fat-fed GNMT KO mice exhibited a decline in total gluconeogenesis (V*_Aldo_*). This was due to diminished gluconeogenesis from phosphoenolpyruvate (V*_Enol_*), but not gluconeogenesis from glycerol (V*_GK_*). In agreement with the decreased gluconeogenesis from phosphoenolpyruvate in 12-week-old, high-fat-fed KO mice, we determined that glucose excursion during a PTT was lower in 32-week-old, high-fat-fed GNMT KO mice compared to WT mice. These gluconeogenic flux characteristics are also consistent with previous reports in low-fat fed mice prior to and in the presence of HCC [11, 17, 20]. Thus, regardless of diet and age, loss of GNMT impedes gluconeogenesis from phosphoenolpyruvate. Importantly, gluconeogenesis is governed by TCA flux and cataplerosis to phosphoenolpyruvate. Total cataplerosis (V*_PCK_*) was lower in GNMT KO mice; however, TCA cycle fluxes (V*_CS_* and V*_SDH_*) were not different between genotypes. This supports the conclusion that the decreased gluconeogenesis from phosphoenolpyruvate was owing to lower partitioning of TCA cycle intermediates towards gluconeogenesis rather than impaired TCA cycle flux.

### 4.2 Integrating SAM homeostasis, metabolic remodeling, and liver pathology in KO mice

The transsulfuration pathway consists of two reactions that result in the synthesis of cysteine. First, CBS generates cystathionine from serine and homocysteine. Second, cystathionine is converted to cysteine by cystathionine γ-lyase (CSE). Prior work has determined that SAM allosterically activates CBS [49, 50], which promotes the entry of homocysteine into the transsulfuration pathway rather than being remethylated to methionine, a SAM precursor. This is consistent with metabolomics analyses in GNMT KO mice presented herein showing the elevated liver SAM is accompanied by lower serine, higher cystathionine, and increased cysteine. Thus, it is reasonable to conclude that elevated SAM stimulates transsulfuration as a means of limiting further SAM accumulation. Of note, transsulfuration is linked to glutathione synthesis [32, 33]. Glutathione is generated from cysteine via two reactions catalyzed by glutamate-cysteine ligase and GSS [32, 33]. In this study, we observed higher GSS protein and glutathione concentrations in KO mice prior to and in the presence of HCC. These results suggest that loss of GNMT stimulates *de novo* glutathione synthesis. This is important from a metabolic perspective because the formation of GSH diverts carbons (glutamate) away from gluconeogenesis. In addition, it has recently been identified that glutathione supports liver lipid accretion by promoting the expression and stability of lipogenic enzymes [51, 52]. Indeed, we identified that ACC, FASN, and SCD1 protein levels were elevated in livers of KO mice. In speculation, loss of GNMT elicits increased transsulfuration and GSH synthesis to mitigate SAM accumulation. This may in turn direct TCA cycle intermediates and precursors away from gluconeogenesis and towards lipid synthesis, desaturation, and accretion.

Another fate of SAM is its use in the synthesis of polyamines. First, SAM undergoes the rate-limiting step of decarboxylation [16]. Decarboxylated SAM then donates an aminopropyl group to form polyamines, leaving MTA as a byproduct [16]. In our study, the redirecting of excess SAM toward polyamine synthesis is reflected by elevated putrescine, spermidine, and MTA in GNMT KO mice. Interestingly, we observed a concurrent increase in liver N^1^-acetylspermidine in GNMT KO mice. SSAT-mediated acetylation of spermidine enables its breakdown and reconversion to putrescine, which sustains polyamine turnover as a catabolic outlet for excess SAM [11, 17, 53]. It may be that the SAM-mediated increase in polyamine synthesis initiates a ‘metabolic pull’ that redirects acetyl-CoA away from oxidation in the TCA cycle for spermidine acetylation so that the liver can better dispose of the elevated SAM in KO mice. Notably, work by others demonstrates that polyamines promote cell growth and proliferation [62, 63], polyamines are higher in HCC [16, 64, 65], and overexpressing SSAT increases susceptibility to tumorigenesis [66, 67]. Together, these findings suggest that the metabolic diversion of acetyl-CoA toward heightened polyamine turnover in GNMT KO mice may contribute to HCC pathophysiology.

### 4.3 SAAR mitigates metabolic remodeling and prevents pathology in livers of KO mice

The results presented in this study support the conclusion that the elevated SAM in GNMT KO mice redirects TCA-cycle intermediates and/or precursors away from gluconeogenesis toward GSH synthesis, lipogenesis, and polyamine turnover. Moreover, these metabolic responses to the loss of GNMT in mice are accompanied by the appearance of fatty liver at 12 weeks of age that transitioned to HCC at 32 weeks of age. To directly test if these metabolic and pathological phenotypes in GNMT KO mice were SAM-dependent, mice were fed a SAAR diet to prevent the accumulation of liver SAM. Dietary SAAR attenuated the increase in liver methionine and SAM in KO mice. Furthermore, the normalization of liver SAM was associated with the absence of liver pathology in KO mice. Specifically, SAAR blocked the development of liver steatosis and fibrosis as indicated by liver TAGs and hydroxyproline levels, respectively. The dietary intervention also abolished a rise in the cell proliferation marker, MCM2, and the appearance of tumor nodules in KO mice, which implies that the development of HCC in KO mice is SAM-dependent. This distinction regarding the *in vivo* mechanisms underlying HCC in the absence of GNMT is important because prior research reports that the tumor suppressor function of GNMT may be independent of SAM levels. Studies in cancer cell lines demonstrated that catalytically inactive GNMT elicited an anti-proliferative effect similar to that of the WT enzyme [35]. Our results do not test the extent to which SAM-independent mechanisms drive liver pathology in KO mice; however, they do suggest that loss of GNMT *in vivo* promotes the development of fatty liver and transition to HCC at least partly via increased SAM availability.

In addition to preventing liver pathology, dietary SAAR mitigated many of the metabolic perturbations observed in GNMT KO mice. SAAR elevated serine, normalized cystathionine, and lowered glutathione concentrations in KO mice. This suggests that the dietary intervention prevented the SAM-driven increase in transsulfuration and glutathione synthesis. Given that glutathione promotes the expression and stability of lipogenic enzymes, the lower glutathione in SAAR-fed KO mice is consistent with the concurrent decrease in ACC, FASN, and SCD1 protein and the reduction in liver TAGs. Moreover, it supports a role for SAM-driven glutathione synthesis in promoting liver steatosis in GNMT KO mice. Reducing SAA intake also increased ornithine, decreased polyamines (putrescine and spermidine), and elevated acetyl-CoA. These results are in agreement with SAAR blocking the rise in polyamine turnover previously linked to tumor initiation and proliferation in GNMT KO mice. The higher acetyl-CoA also supports the hypothesis that polyamine acetylation uses a shared substrate pool with the TCA cycle and that this competition for acetyl-CoA is diminished when SAM accumulation is prevented.

Notably, SAAR did not rescue KO mice from displaying reduced glucose excursion during a PTT, a surrogate for gluconeogenesis. This contrasts with our previous findings in low-fat-fed mice wherein SAAR increased TCA cycle cataplerosis and gluconeogenesis from phosphoenolpyruvate [20]. This discrepancy indicates that the ability of SAAR to restore gluconeogenesis may be diet-dependent. It is also possible that the PTT may not fully reflect whether SAAR resolves the impaired gluconeogenesis resulting from loss of GNMT. Our prior work showed that SAAR independently enhances *de novo* serine synthesis [68], which could divert pyruvate-derived gluconeogenic intermediates away from glucose production. Thus, future work using isotope-based flux approaches is required to disentangle the effects of SAAR on gluconeogenesis from its effects on serine synthesis in GNMT KO mice.

In conclusion, loss of GNMT in diet-induced obese mice redirects liver TCA cycle intermediates and/or precursors away from gluconeogenesis toward SAM-dependent biosynthetic pathways. This is accompanied by the development of fatty liver and progression to HCC. Dietary SAAR prevents SAM accumulation, mitigates the metabolic remodeling, and attenuates the appearance of liver pathology. These findings support that GNMT insufficiency promotes MASLD-related HCC via SAM-dependent mechanisms.

## Data Availability

Data will be made available upon request.

## CRediT Authorship Contribution Statement

**Griffin S. Hampton:** Investigation, Formal analysis, Data curation, and Writing - review & editing. **Andres F. Ortega:** Investigation, Formal analysis, Data curation, and Writing - review & editing. **Cha Mee Vang:** Investigation, Formal analysis, Data curation, and Writing - review & editing. **Ferrol I. Rome:** Investigation, Data curation, and Writing - review & editing. **Mickael Goelzer:** Investigation and Writing - review & editing. **Louise Lantier:** Investigation, Formal analysis, Data curation, Resources, and Writing - review & editing. **Curtis C. Hughey:** Conceptualization, Investigation, Formal analysis, Data curation, Resources, Funding acquisition, Supervision, Project administration, Visualization, Writing – original draft, and Writing – review & editing.

## Acknowledgements

We thank Ruth Pfeiffer, Jared Hartmann, and Deanna Bracy for their technical assistance in methodological procedures. The authors acknowledge the technical support provided by the Vanderbilt Mouse Metabolic Phenotyping Center Body Weight Regulation, Analytical Resources, and Surgical Services Cores. Figure schematics were generated using BioRender.com.

---

**Figure S1. Loss of glycine N-methyltransferase (GNMT) increases energy expenditure and lowers adiposity in mice.** Twelve-week-old wild type (WT) and GNMT KO mice were a high-fat diet for 6 weeks prior to body composition measurements and indirect calorimetry experiments. **A:** Body weight (g), fat (g), and lean mass (g) in WT and KO mice (n = 8-13 per genotype). **B:** Energy expenditure (kcal·hr^-1^; n = 8-13 per genotype) for 12-hour light and dark cycles. **C:** Energy expenditure (kcal·hr^-1^; n = 8-13 per genotype) for 24 hours. **D:** Respiratory exchange ratio (RER; unitless; n = 8-13 per genotype) for 12-hour light and dark cycles. **E:** RER (unitless; n = 8-13 per genotype) for 24 hours. **F:** Food intake (g; n = 8-13 per genotype) for 12-hour light and dark cycles. **G:** Food intake (g; n = 8-13 per genotype) for 24 hours. **H:** Locomotor activity by mice (All meters; n = 8-13 per genotype) during 12-hour light and dark cycles. **I:** All meters by mice (n = 8-13 per genotype) for 24 hours. Data are mean ± SEM. Statistical differences were determined by a Student’s t test and accepted as significant if p<0.05 for data in panel A. Statistical differences (p<0.05) for data in panels B-I were determined by a two-way ANOVA. Significant main effects are presented within each panel. **p<0.01. @ p<0.05 by ANCOVA when regression is run against body weight.

