## Supplemental Table 2 for "Sulfur amino acid restriction prevents S-adenosylmethionine-driven liver steatosis, hepatocellular carcinoma, and metabolic remodeling in high-fat-fed GNMT-null mice"

| Antibody | Supplier | Catalog Number | RRID | Dilution |
|---|---|---|---|---|
| Acetyl-CoA carboxylase | Cell Signaling Technologies | 3662 | RRID:AB_2219400 | 1:1000 |
| ATP-citrate lyase | Cell Signaling Technologies | 4332 | RRID:AB_2223744 | 1:1000 |
| Cystathionine β-synthase | Proteintech | 14787-1-AP | RRID:AB_2070970 | 1:1000 |
| Gamma Cystathionase | Proteintech | 12217-1-AP | RRID:AB_2087497 | 1:1000 |
| Glutamate-cysteine ligase catalytic subunit | Proteintech | 12601-1-AP | RRID:AB_2278734 | 1:1000 |
| Glutathione synthetase | Proteintech | 15712-1-AP | RRID:AB_2878171 | 1:1000 |
| Glycine N-methyltransferase | Proteintech | 18790-1-AP | RRID:AB_10597093 | 1:1000 |
| Fatty acid synthase | Cell Signaling Technologies | 3180 | RRID:AB_2100796 | 1:1000 |
| Minichromosome maintenance 2 | Cell Signaling Technologies | 4007 | RRID:AB_2142134 | 1:1000 |
| Spermidine/spermine N1-acetyltransferase 1 | Proteintech | 10708-1-AP | RRID:AB_2877739 | 1:1000 |
| Stearoyl-CoA desaturase 1 | Cell Signaling Technologies | 2794 | RRID:AB_2183099 | 1:1000 |
| Anti-rabbit IgG | Cell Signaling Technologies | 7074 | RRID:AB_2099233 | 1:5000 |

**Supplementary Table 2. List of antibodies.**
