## Supplementary figures and images for "Sulfur amino acid restriction prevents S-adenosylmethionine-driven liver steatosis, hepatocellular carcinoma, and metabolic remodeling in high-fat-fed GNMT-null mice"

### Supplemental Figure 1

**FIGURE S1. Lower adiposity is associated with higher energy expenditure in GNMT KO mice**

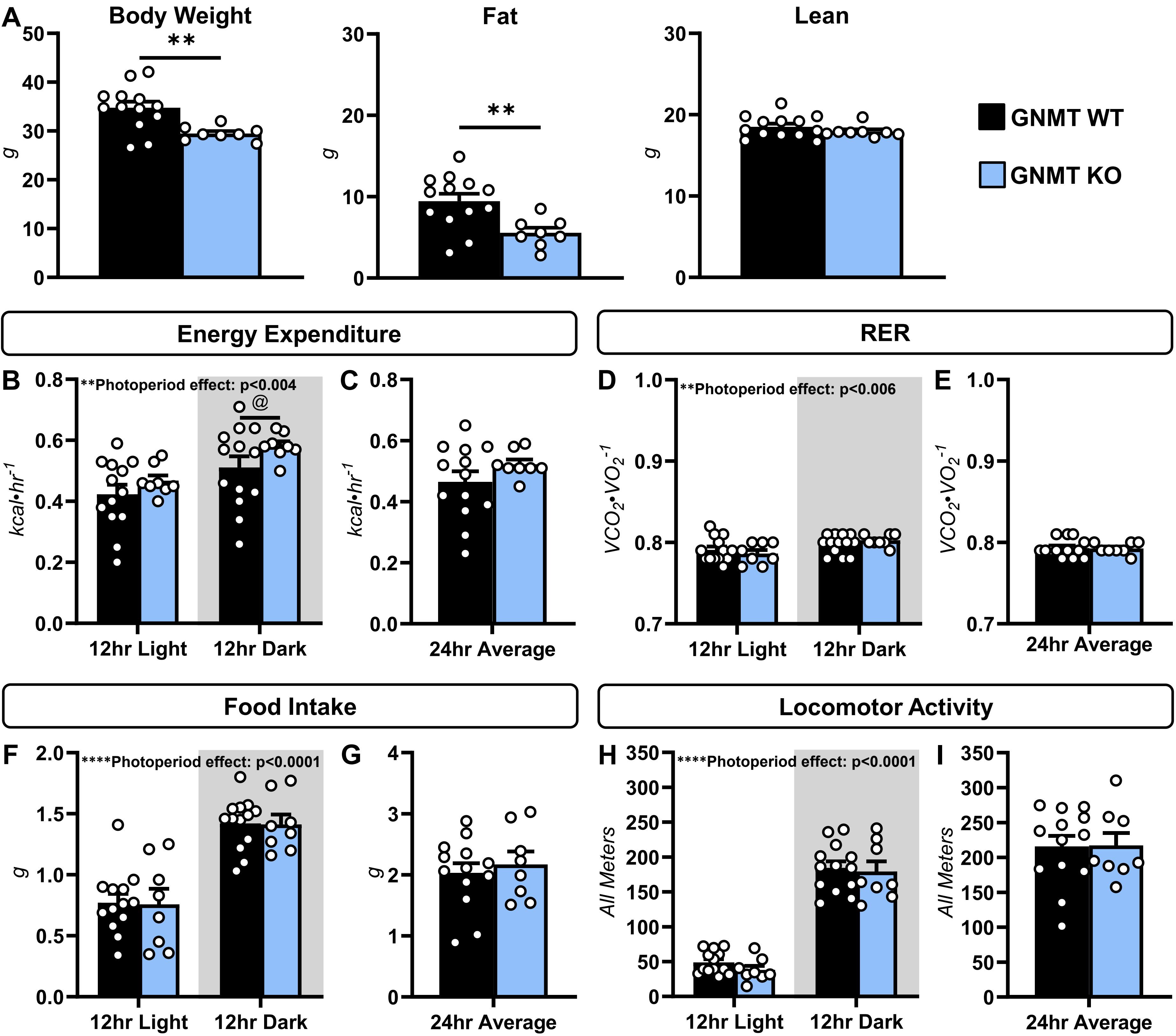
